# The algorithmic neuroanatomy of action-outcome learning

**DOI:** 10.1101/137851

**Authors:** Richard W. Morris, Amir Dezfouli, Kristi R Griffiths, Mike E Le Pelley, Bernard W Balleine

**Author notes:** Correspondence: Bernard Balleine Decision Neuroscience Laboratory Level 4 Matthews Building School of Psychology University of NSW Kensington, NSW 2052 Australia.

## Abstract

Although it is well known that animals can encode the consequences of their actions and can use this information to control action selection and evaluation, it is not known what learning rules control action-outcome (AO) learning. Here we trained participants to encode specific AO associations whilst undergoing functional imaging (fMRI) and used computational modelling to evaluate competing models. This analysis revealed that a Kalman filter, which learned the unique causal effect of each action, best characterized AO learning and found the medial prefrontal cortex differentiated the unique effect of actions from background effects. We subsequently extended these findings to show that mPFC participated in a circuit with parietal cortex and caudate nucleus to segregate distinct contributions to AO learning. The results extend our understanding of goal-directed learning and demonstrate that sensitivity to the causal relationship between actions and outcomes guides goal-directed learning rather than contiguous state-action relations.

---

There is now considerable evidence that animals are capable of encoding the consequences of their actions and that they use that information to select, evaluate and initiate future actions.^1–5^ Although it is clear, therefore, that such learning involves encoding of the action-outcome relationship, the learning rules that govern that learning have yet to be established. Normatively, this relationship has been described by the formalism ΔP (Figure 1A); which captures the effect of the action on outcome delivery over and above any background effects^6,7^. Nevertheless, how we distinguish these effects is unclear.

**Figure 1.**
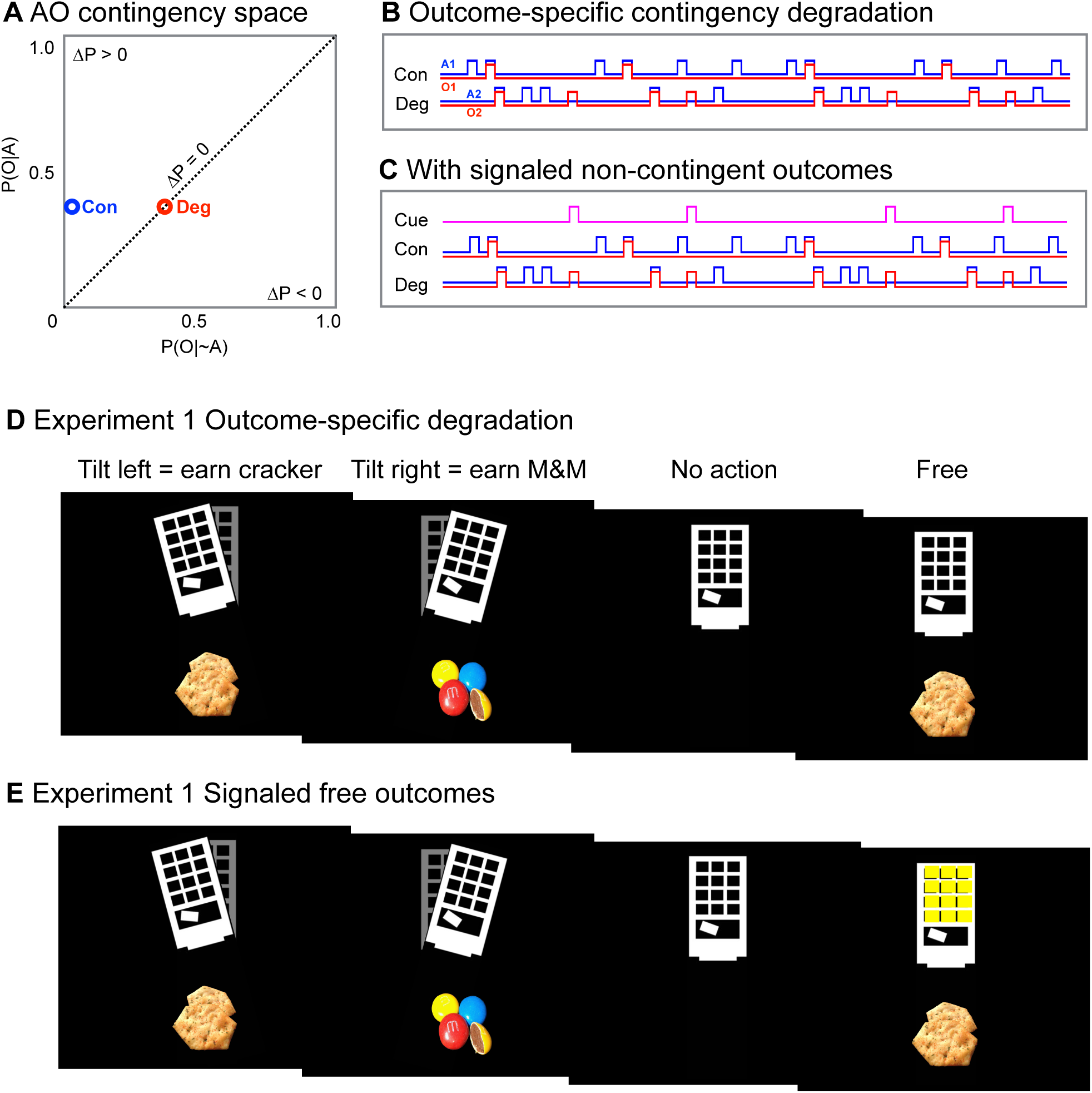
The action-outcome relationship in **A)** contingency space, where ΔP = P(O|A) – P(O| ~ A), that is, a positive ΔP exists when the conditional probability of an outcome given an action is greater than the probability of the outcome in the absence of the action (Con) while ΔP approaches zero as these conditional probabilities become equal (e.g., Deg); **B)** Outcome-specific degradation schedule where P(O1|A1) = P(O2|A2) while the addition of noncontingent outcomes (O2) produces differences in ΔP (i.e., ΔP = 0.4 and 0 for Con and Deg, respectively); **C)** Signaled schedule where the cue (pink) marks the noncontingent outcomes **D)** The action-outcome relationships presented onscreen (counterbalanced) in the degradation test in the MRI and **E)** the signaled follow-up test after the MRI, where a 1-s visual cue (yellow) indicated the delivery of each noncontingent outcome

There are, in fact, very few mechanistic models of how we and other animals detect and encode the AO contingency. Historically, models of associative learning have been proposed as an algorithmic account of causal learning, including learning about actions.^8,9^ These models, and their modern varients,^1^ assume that background conditioning plays a key role because the background competes with actions via a summed error term, to uniquely predict the outcome. More recently model-based reinforcement learning models (MBRL), which forgo a summed error-term and instead incorporate a model of the task structure, have been proposed as a general account of goal-directed action.^4,10,11^ MBRL represents the task structure in a covariance matrix that represents the contiguity, rather than the unique causal relationship, between actions and outcomes (i.e., the state-action transitions).

Here we evaluated the neural computations of learning about causal actions (AO learning) in the human brain. We found a simple Kalman filter^12–14^ that combined prediction-error learning with the covariance structure of the environment explained the acquisition of AO learning better than models based on covariance or prediction-error alone. This iterative model attributed prediction-errors to different causal variables adjusted by their covariance, in order to uniquely predict the outcome. Critically, model-based fMRI revealed activity in the medial prefrontal cortex (mPFC) and the dorsal anterior cingulate cortex (dACC) tracked changes in the predictive value of actions and the background with respect to specific outcomes, separately. Furthermore, we found the mPFC participated in a network with the striatum and posterior parietal cortex to segregate the influence of different causes via their covariance, a unique prediction of the Kalman algorithm. These findings reveal, for the first time, an integrated corticostriatal network that combines prediction-errors with covariance in order to learn how our actions control our environment.

## Results

We performed two fMRI experiments to probe the computational mechanisms by which the brain learns the AO contingency and distinguishes it from any background effects. In typical neuroscience experiments, actions and their consequences are offered in discrete trials where there is no ambiguity about unique effects. Instead we used a free-operant task without a discrete trial structure, along with noncontingent outcomes, which required participants to infer the causal effects of their actions.

### AO contingency degradation revealed people learned the unique effects of their actions

Experiment 1 involved training hungry participants with two actions for distinct food outcomes, selected before the task by each participant (e.g., button 1 = M&Ms, button 2 = BBQ flavored crackers). Training occurred according to a ‘constant probability’ schedule.^15^ During the fMRI test, at the end of every second the schedule recorded whether a button was pressed and then presented an outcome onscreen with a conditional probability P(Oi|Ai) = 0.2, for each action. (Actual food outcomes were provided at the end of all testing). In order to selectively degrade the causal relationship of one AO contingency while equating the reward value of both actions, the schedule also presented one of the outcomes onscreen if neither button had been pressed, i.e., P(O1| ~A1, ~A2) = 0.2. Under this outcome-specific degradation schedule, delivery of the noncontingent outcome diminishes the reward value of both actions equally (since reward can now be obtained without taking either action). However the noncontingent outcome will selectively reduce the *causal* relationship of only one action (A1) and not the other (A2), because the noncontingent outcome is indistinguishable from the outcomes caused by one action, but easily distinguishable from the outcomes caused by the other action (see methods).^16^

Figure 2A illustrates that a preference for the non-degraded action (Con) clearly emerged over time as people were exposed to the differences between each AO contingency. Figure 2B (left panel) shows that overall the mean number of Con actions was greater than the degraded actions (Deg). Causal ratings collected at the end of each two minute block also confirmed people judged the Con action more causal than the Deg action, shown in Figure 2C (left panel).

**Figure 2.**
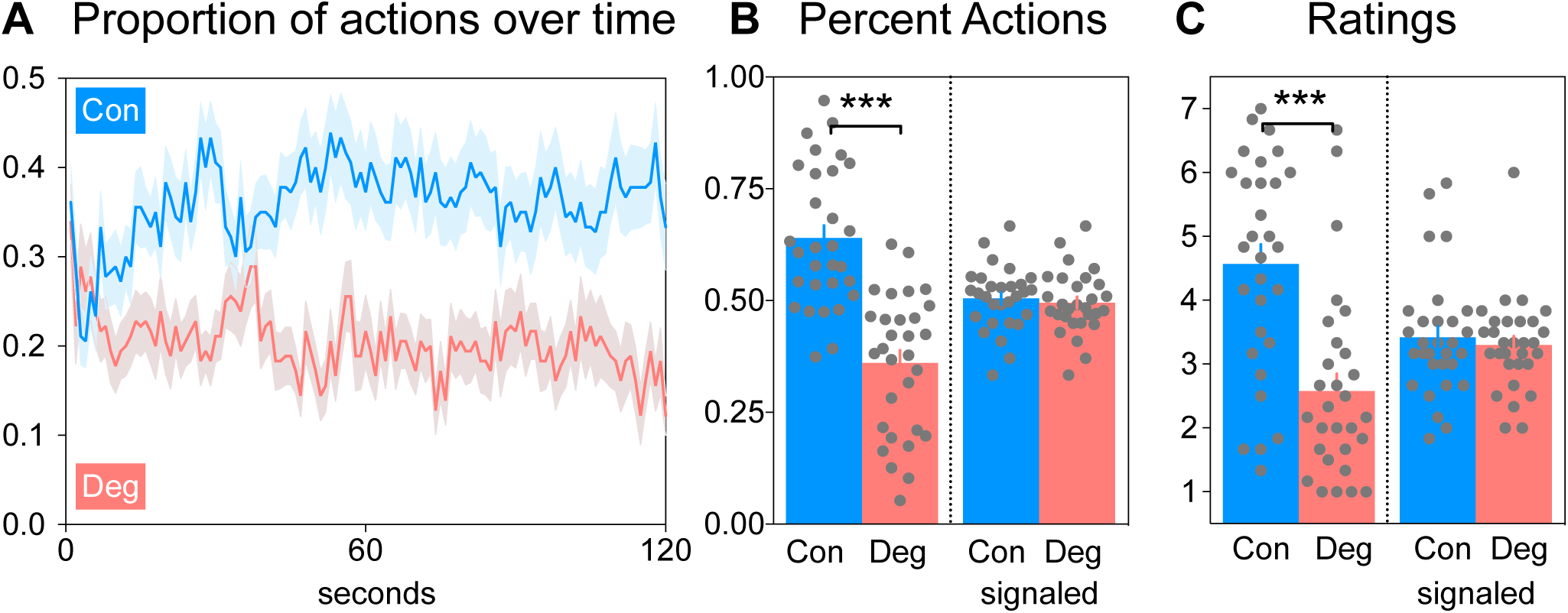
Experiment 1 behavioural results (*N* = 30) **A)** Mean (shaded = SEM) probability of each action over time shows that on average the contingent action was gradually selected over the degraded action **B)** The mean (errorbars SEM) percent of contingent actions was significantly greater than degraded actions when free outcomes were unsignalled, paired t-test *t*_29_ = 4.15, \*\*\**p* = .0002. However when free outcomes were signalled then degraded actions were restored, paired t-test *t*_29_ = 0.75, *p* = .46. **C)** Mean (errorbars SEM) causal judgments of the contingent action were greater than the degraded action when free outcomes were unsignalled, paired t-test *t*_29_ = 3.94, \*\*\**p* < .0004, and this difference was removed when the free outcomes were signalled, paired t-test *t*_29_ = 0.88, *p* = .39.

The selective impact of noncontingent outcomes on actions and causal judgments reveals our sensitivity to the unique effect of our actions, even when the probability of reinforcement among actions is equal. The noncontingent outcomes made it more difficult for participants to distinguish the outcome they caused from the outcome that would have occurred anyway. As a result, the perceived causal efficacy of that action was reduced. We conducted a follow-up test after the fMRI scan, under the same contingencies, with the addition that each noncontingent outcome was now signaled by a yellow-light cue (Figure 1B & D). The signal reduces the uncertainty about the noncontingent outcomes, which allows participants to once again distinguish the unique effect of their own actions. The results found the addition of the signal restored responding (Figure 2B, Signaled) as well as causal judgments of the degraded action (Deg) to the same level as the Con action (Figure 2C, Signaled). The restoration of actions and causal judgments by the signal implies that learning about the base-rate or ‘background conditioning’ plays a key role in learning the unique causal strength of our actions.

### Action-selection reflected causal learning rather than reinforcement learning

Figure 3A shows the correlation between causal judgments and causal actions for each person in Experiment 1 was high and significant, consistent with causal learning guiding action-selection rather than the frequency of reinforcement or immediate temporal effect of reward. To check our experimental control over each contingency in this free-response task, we confirmed the noncontingent outcomes selectively degraded the experienced contingency of the degraded action: post-hoc analysis revealed the mean contingencies experienced for the Con and Deg action were ΔP = 0.18 and ΔP = 0.07 respectively, paired t-test *t*_29_ = 12.06, *p* = .8E-12. The positive contingency (ignoring noncontingent outcomes) for the Deg action was P(O1|A1) = 0.17, and very similar to the Con action (P(O2|A2) = 0.18, paired t-test *t*_29_ = 0.90, *p* = .37), confirming that serendipitous differences in the positive contingency were not responsible for the results. Importantly, each person received noncontingent outcomes in each block, with the exception of one person who received noncontingent outcomes in only 4 out of 6 blocks. We also checked whether any serendipitous reward contingency existed for the Con action. Figure 3B (blue) shows the correlation between the number of Con actions and the total number of outcomes (contingent + noncontingent) delivered was close to zero across participants, confirming there was no serendipitous reward contingency that may have influenced preference for the Con action. Conversely there was no negative contingency between Deg actions and total outcomes (Figure 3B, red). Furthermore, the distribution of delays between each outcome and the preceding action did not differ within a 10-s interval for Con and Deg actions (Figure 3C), confirming that the immediate temporal contiguity with reward was not differentially influencing action-selection. Finally, pre-test preference ratings of the snack food outcomes confirmed both rewards were equally liked. The mean (95% confidence interval) ratings on a 7-point Likert scale were 5.8 CI[5.5, 6.2] and 6.3 CI[5.9, 6.6], for BBQ crackers and M&M respectively. Thus, action-selection did not reflect any post hoc or serendipitous differences in reward contingency or contiguity.

**Figure 3.**
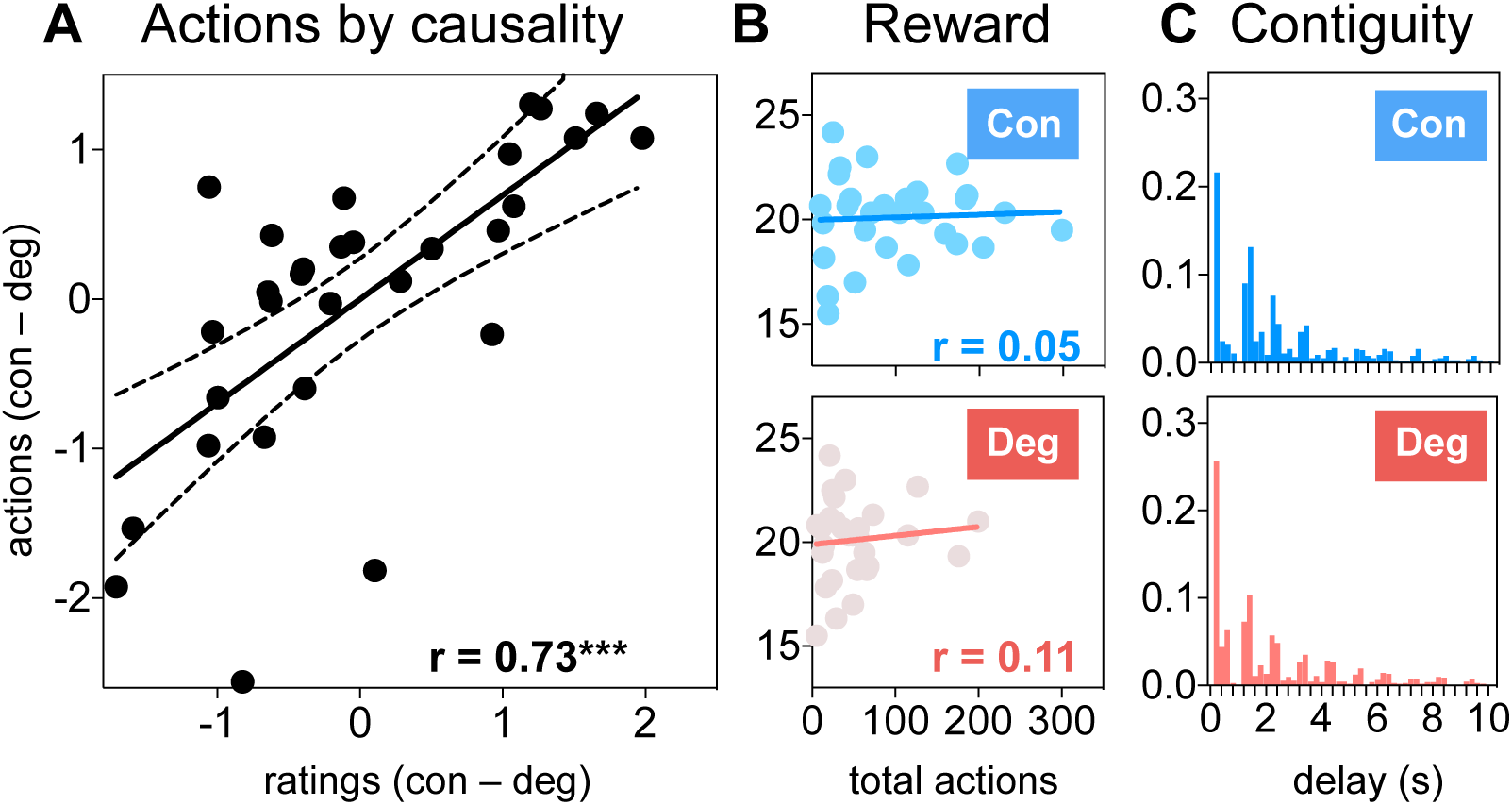
Causality explains action selection better than reward in Experiment 1 (*N* = 30) **A)** The difference (in z-score units) in contingent over degraded actions and causal judgments among participants was correlated, *F*_1,28_ = 26.20, \*\*\**p* = .00002 (dotted lines 95% CI). **B)** No correlation existed between the number of contingent actions (blue, *F*_1,28_ = 0.07, *p* = .79), or degraded actions (red, *F*_1,28_ = 0.30, *p* = .59) and the total number of outcomes among participants. **C)** Frequency histogram of the experienced delays between actions and reward shows the distribution was similar for both actions, Kolmogorov-Smirnov *D*_78_ = 0.09, *p* = 0.99

### Modelling revealed a Bayesian prediction-error best explained AO learning

We simulated and fit three models: a prediction-error model with a summed prediction-error term, a MBRL model with a covariance matrix, and a Kalman algorithm with both, to determine which best explained behavioral performance in Experiment 1. The prediction-error model assumed actions and background cues competed for causal strength via a summed error-term (see methods). Simulation (Supplementary figure 1) confirmed this model resolved the unique effect of the causal action (i.e., converged to ΔP). By contrast, the MBRL used a transition matrix (updated via a state prediction-error) to represent the covariance between each action, outcome and background. Simulation confirmed the covariance learned by this model was insufficient to distinguish the causal action. In particular, the reward value of the free outcomes outcompeted both actions equally in the MBRL, which did not learn or prefer the causal action. Finally, the Kalman algorithm assumed actions compete with background effects via a summed prediction-error term, however the amount learned about each is adjusted by the covariance between them. That is, when causal variables covary (i.e., positive covariance), the model cannot distinguish their separate influence and so adjusts them together. However with negative covariance then the effects can be distinguished and changes in the belief of one cause will inversely affect the other (see Figure 4). By combining the prediction-error term with the covariance, the Kalman filter distinguished causal actions across the widest range of parameter values (Supplementary figure 2).

**Figure 4.**
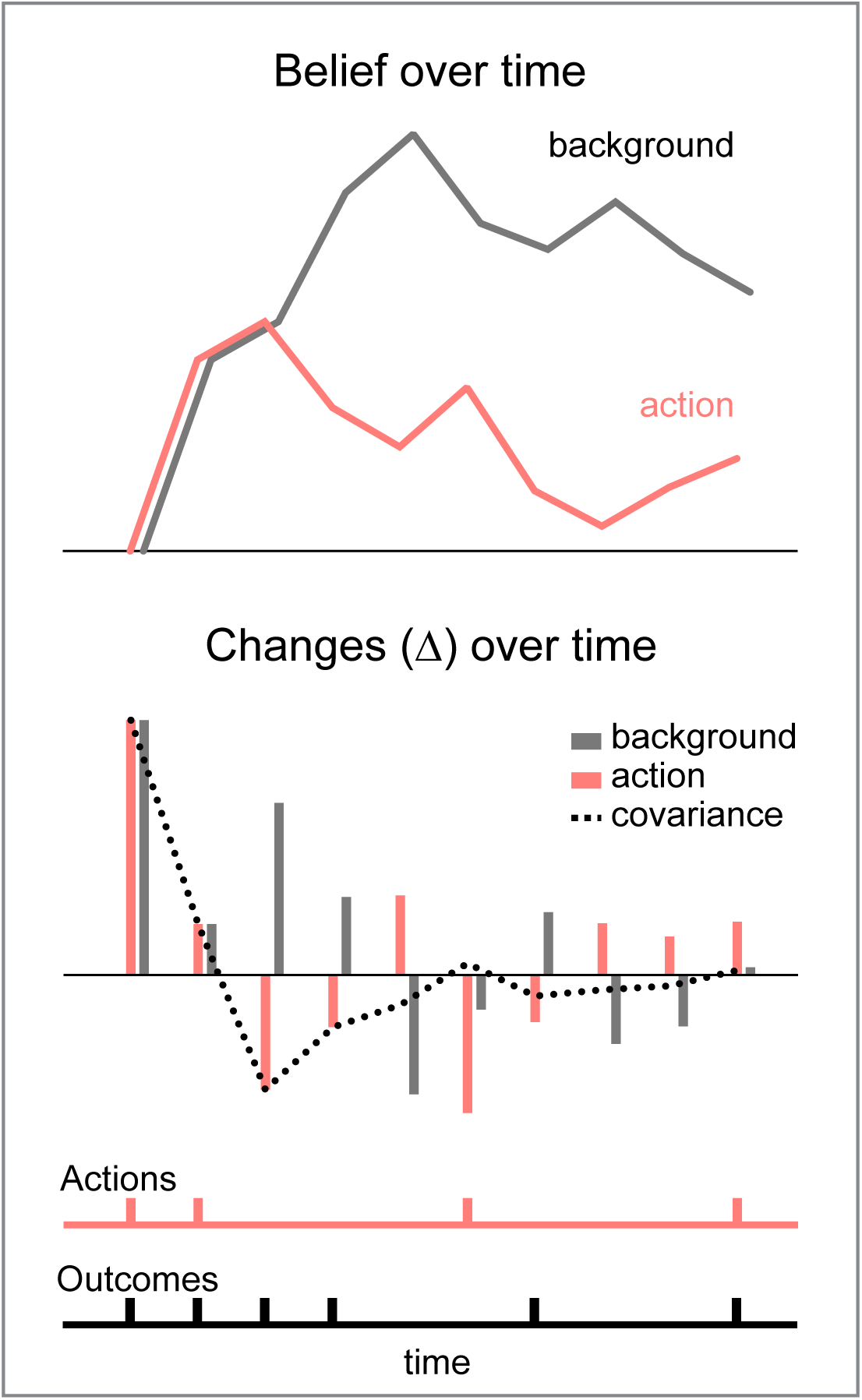
The Kalman algorithm changes beliefs over time according to a prediction-error adjusted for covariance **A)** Under a degradation schedule, belief in the action and background initially increase together as actions co-occur with outcomes. However with free outcomes, the belief in the background diverges from the action. **B)** Changes in the background and action occur in the same direction when covariance between the action and background is positive, but when the covariance is negative then beliefs move in opposite directions. **C)** Action and outcome events over time in this example

Behavioral fitting indicated action selection was better explained by the Kalman algorithm than a prediction-error model or MBRL. Data from Experiment 1 was used to calculate the posterior probability of the Kalman algorithm, as well as a prediction-error model, a MBRL model (using a transition matrix), and a null model. The null model used the asymptote AO contingencies of each block as action values, thus it was not a learning model but a static model with no temporal dynamics. Comparison with the null model determines whether each learning model can explain how choices depend on the sequence of trial-by-trial feedback. Table 1 shows the negative log likelihoods and relative Bayes factors of each model relative to the null. After fitting each participants’ data by maximum-likelihood estimation, the results of a likelihood ratio test (LRT) indicated all learning models predicted significantly more behavior than chance. However, the relative Bayes factor (vs the null model) shows only the optimal Kalman algorithm had a positive value, indicating only this model predicted the acquisition of causal learning over time. This model also explained more choices and the behavior of more individuals than the other models, with a Pseudo-R2 of 0.24 and a positive evidence ratio of 2 (20/10 favoring H_1_/H_0_), which were higher than the respective values for the other learning models. Thus, the majority of subjects and the total evidence favors the simple Kalman algorithm over a static model with no temporal dynamics but perfect asymptote performance. This was not the case for the optimal MBRL, or the prediction-error model (or a causal induction model, see Supplementary Materials), which all explained more variance than chance but had a negative GBF relative to the null and a PER less than 1 (Table 1).

**Table 1.** Model evidence and comparison scores

| | Free parameters | Negative log likelihood | LRT $X^2$ | LRT $p$ | Pseudo-$R^2$ | Relative Bayes Factor ($H_1 - H_0$) | No. favoring $H_1/H_0$ model |
| --- | --- | --- | --- | --- | --- | --- | --- |
| Kalman filter | $\nu$, $k$, and $\tau$ | 14,056 | $X^2_3 = 7,790$ | $< 1E-30$ | 0.24 | 1573 | 22/6 |
| Prediction-error | $a$, $k$, and $\tau$ | 16,343 | $X^2_1 = 5,124$ | $< 1E-30$ | 0.13 | -506 | 18/11 |
| MBRL | $a$, $k$, and $\tau$ | 18,853 | $X^2_1 = 1,04$ | .7E-15 | 0.01 | -2959 | 1/25 |
| Null model ($H_0$) | $\tau$ | 15,992 | $X^2_1 = 5,826$ | $< 1E-30$ | 0.13 | | |

Legend: For each model, optimal model evidence (aggregate negative log likelihood scores), model significant differences from chance (Likelihood ratio test, LRT), model fit (Psuedo- $R^2$ ), as well as Bayesian model comparisons among each alternate model relative to the informed model ( $H_0$ ). $\nu$ is prediction uncertainty, $\alpha$ is learning rate, $k$ is the AO delay temporal threshold, and $\tau$ is inverse temperature (exploitation/exploration).

### Model-based fMRI revealed the mPFC distinguishes causal actions from background effects

We evaluated whether the brain learned about causal actions, as described by the Kalman filter, by regressing the model-derived learning estimates against the image data collected in Experiment 1. Independent learning estimates for actions and background (ΔAO and ΔXO; see Methods) were included as parametric modulators of a stick (delta) function of response and outcome times. We included outcomes in the delta function in order to include times when the action was present as well as absent (the background was assumed to be always present). Whole-brain analysis revealed learning about actions and the background occurred in distinct regions of the mPFC (Figure 5A & B). Action learning (ΔAO) appeared in a medial region of the superior frontal gyrus (BA9, global peak MNI co-ordinates: -15 +47 +40, Z = 4.71, *F*_1,29_ = 37.12, FWE = .031). At the same time, learning related changes to the background estimates (ΔXO) appeared in the dorsal anterior cingulate cortex (BA32, global peak MNI: -9 +41 +22, Z = 5.19, *F*_1,29_ = 50.20, FWE = .004), as well as smaller changes in the left caudate (MNI: -15 +14 +7, Z = 4.88, *F*_*1,29*_ = 41.80, FWE = .017), and cuneus (MNI: -3 -64 +34, Z = 4.39, *F*_*1,29*_ = 37.28, FWE = .04). No other region survived multiple comparison correction in this whole-brain analysis.

**Figure 5.**
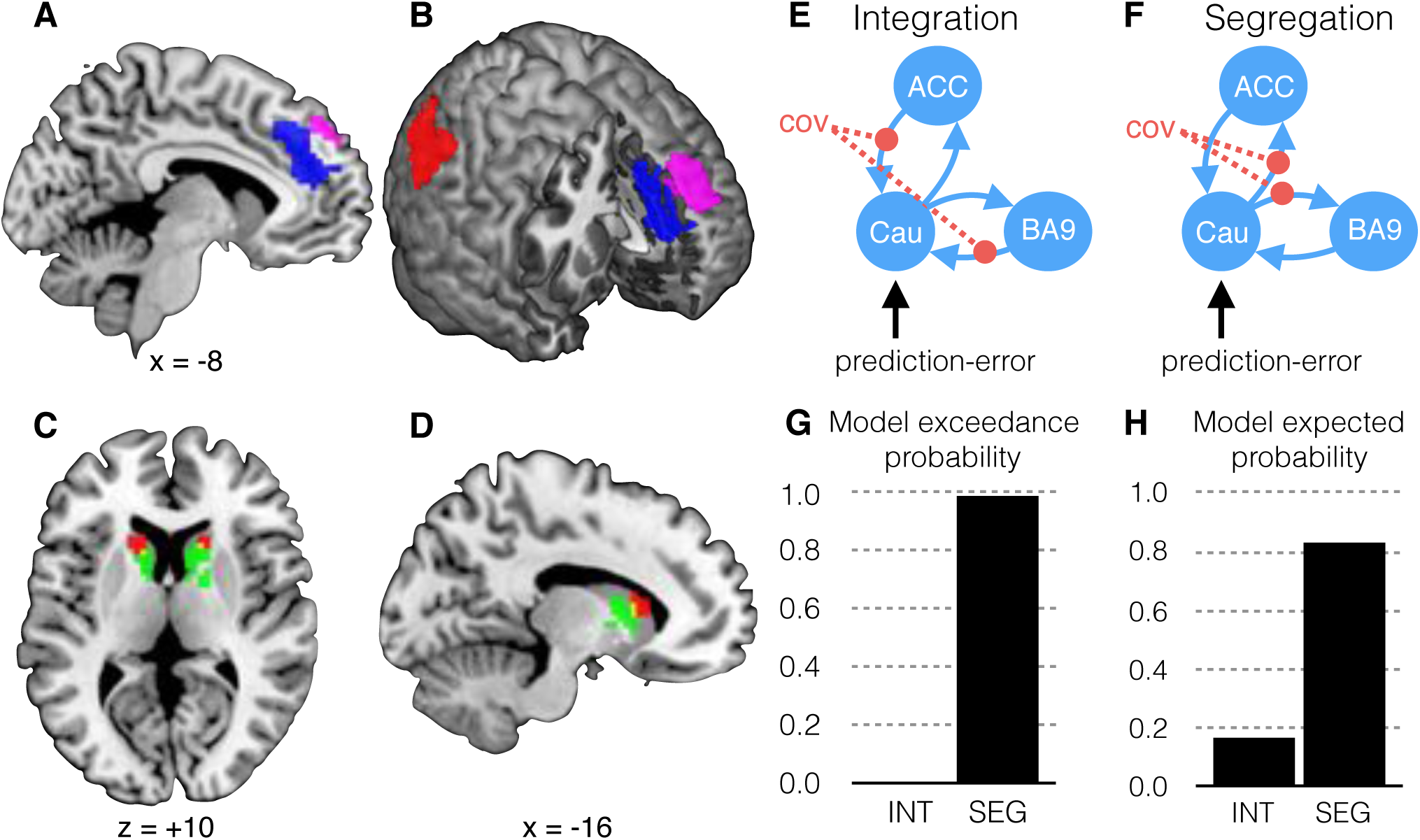
Corticostriatal network for causal learning in Experiment 1, *N* = 30. **A**) Model-derived learning variables were tracked in the medial prefrontal cortex: Model updates to actions (*Δ*AO) occurred in the medial prefrontal cortex (BA9) (violet voxels FWE cluster *p* = .007). At the same time, model updates to the background (*Δ*XO) occurred in the dorsal anterior cingulate cortex (ACC), (blue voxels FWE cluster < .001); **B**) Cut-away representation showing the spatial relationship of the corticostriatal network, including model covariance in the right posterior parietal cortex (BA40) (red voxels FWE cluster *p* =). **C**) ROI analysis in the striatum: Red voxels (image threshold *p* < .05 svc) in the anterior caudate tracked the covariance, while green voxels (image threshold *p* < .05 svc) in the caudate body tracked summed prediction-errors. Overlap is indicted in yellow. **D**) Sagittal view of ROI results. **E**) DCM showing the caudate integrates information from separate regions in the mPFC (dACC and BA9), modulated by the covariance between potential causes; **F**) An alternate DCM showing the caudate segregating information in the mPFC, modulated by the covariance between potential causes **G**) The probability the data supports the Segregate model (SEG) is more likely than the Integrate model (INT). **H**) Posterior probability of each model (INT vs SEG) generating the observed data

The results described so far indicate the background expectancy produced by the noncontingent outcome plays a key role in learning the unique effect of our actions. According to prediction-error models (including the Kalman algorithm), this background expectancy will be violated whenever the noncontingent outcome does not follow an action. This implies that a negative AO contingency will be learned under certain conditions (i.e., inhibitory learning). For example, in Experiment 1 we explicitly arranged that O1 sometimes occurs after A1 but never after A2 (in order to equate the reward value of both actions), which results in a negative contingency between A2-O1. We tested a regressor of changes to this negative AO contingency, as learned by the Kalman algorithm, in the whole-brain. BOLD activity in a ventral medial prefrontal region, including the anterior cingulate (BA32) and medial orbitofrontal cortex (BA10), learned the negative AO contingency, global peak MNI: -3 +50 -11, Z = 3.52, *F*_*1,29*_ = 18.08, FWE = .011 (Supplementary figure 3). These imaging results are consistent with recent reports in rodents that the medial orbitofrontal cortex is critical for learning about unobserved outcomes, and in particular a negative AO contingency.^17^

### Model-based fMRI showed the posterior parietal cortex tracks covariance between causes

The results so far indicate causal actions are distinguished from the background in the mPFC. A unique feature of the Kalman filter is that the covariance term distinguishes the influence of candidate causes by updating ΔAO and ΔXO together when the covariance is positive and updating them in opposite directions when the covariance is negative (Figure 4). We tested whether any brain regions tracked the covariance between actions and background by entering the covariance values as a parametric modulator. A whole-brain analysis revealed bilateral activity in posterior parietal cortex (BA40) was significantly associated with the covariance term, left global peak MNI: -57 -55 +37, *Z = 5.83, F*_*1,29*_ = 72.78, FWE < .001 and right peak MNI: +42 -67 +43, Z = 5.34, *F*_*1,29*_ = 54.67, FWE = .002 (Figure 5B).

### ROI analysis showed prediction-error and covariance converge in caudate

Substantial evidence exists that the ventral striatum tracks or receives reward prediction-errors,^18–20^ while dorsal striatal regions track action values.^18^ In the present example of AO learning, the prediction-error represents the deviations between the observed outcome and the summed total causal expectancy (action + background). We tested whether the striatum tracks this summed error term, by including it as a parametric modulator in an anatomical ROI analysis of the striatum. Figure 5C & 5D shows BOLD responses in a posterior region of the caudate body (green) tracked the summed errors (ROI peak MNI:: +15 +11 +4, *Z* = 4.17, *t*_29_ = 4.94, FWE = .002 svc) while activity in the anterior caudate (red) was associated with the covariance between actions and background (ROI peak MNI: -15 +23 +7, *Z* = 3.42, *t*_29_ = 3.84, FWE = .029 svc) These regions were more medial and dorsal than those implicated in reward prediction-error signals but similar to regions implicated in instrumental learning.^18^ Thus, the caudate appears to receive sufficient information to segregate the influence of different events and may play an important role in selectively distinguishing causal actions from background effects.

### DCM revealed the caudate segregates the effect of the action from background effects

To further determine the caudate’s role in distinguishing control, we performed a dynamic causal model (DCM) analysis.^21^ We tested two possibilities shown in Figure 5E & 5F. In Model 1 the caudate is a site of convergence of the updated values from the prefrontal cortex to enable action-selection. In Model 2 the caudate segregates the prediction-error to update the estimates of the action and background separately. Bayesian model selection revealed the relative log-evidence for model 2 was 85.31, which corresponds to strong evidence in favor of segregation. A random-effects analysis (Figure 5G & 5H) revealed a large majority of participants were significantly more consistent with the segregate model than the integration model (the exceedance probability was 99 percent), and it was more likely to be true for any random subject (the posterior probability of the segregate model was 80 percent).

### PPI showed cortex and caudate interact when causal actions must be distinguished by their covariance

Experiment 2 replicated the key fMRI results in an independent sample of naive participants, using a design that allowed us to assess the effect of distinguishing the free outcomes on the corticostriatal network we identified above. The experiment used a single AO contingency and varied whether or not the free outcomes were distinguishable from the earned outcomes in a block-by-block fashion, to test the interaction between causality and caudate activity. For half the blocks we used the same outcome for both free and earned outcomes (i.e., as in Experiment 1), while in the other half the earned outcomes were different from the free outcomes. Using distinct outcomes in half the blocks allowed the participant to discern the causal effect of their actions (i.e., equivalent to signaling the free outcomes, as in the follow-up to Experiment 1). Figure 6A shows that degradation reduced total actions and causal ratings, however providing distinct free outcomes restored causal actions and judgments. As before, we fitted the Kalman algorithm to the data using maximum likelihood estimation. The optimal model predicted significantly more choices than chance, the mean group average likelihood per trial was 57 percent (95% CI: 53-60). A functional ROI (fROI) analysis using masks generated from the significant results in the mPFC and caudate of Experiment 1 confirmed that learning about the background (ΔXO) occurred in the same dorsal ACC region (Figure 6C, blue), ROI peak MNI: -3 +36 +38, *Z* = 3.94, *t*_19_ = 5.06, FWE = .02. Meanwhile learning values for the action (ΔAO) occurred in the same region of BA9 (Figure 6C, violet), ROI peak MNI: -2 +47 +46, *Z* = 3.04, *t*_19_ = 3.56, *p* = .001. Covariance between the action and background was tracked in the caudate (Figure 6C, right), ROI peak MNI: -12 +20 +1, *Z* = 3.91, *t*_19_ = 5.01, FWE = .002. We used a whole-brain PPI analysis to determine whether any cortical regions interacted with the caudate when free outcomes were indistinguishable from earned outcomes. A single region in the right parietal junction interacted with the caudate when free outcomes were the same (versus different), shown in Figure 6E, global peak MNI: +54 - 58 +30, *Z* = 4.68, *F*_*1,19*_ = 24.90, FWE = .01. This region overlapped with the right posterior parietal cortex (BA40) identified in Experiment 1 (Figure 6E, red). Together these results implicate the posterior parietal cortex in tracking the covariance with the background, and interacting with the caudate when the unique effect of our actions must be distinguished from the background.

**Figure 6.**
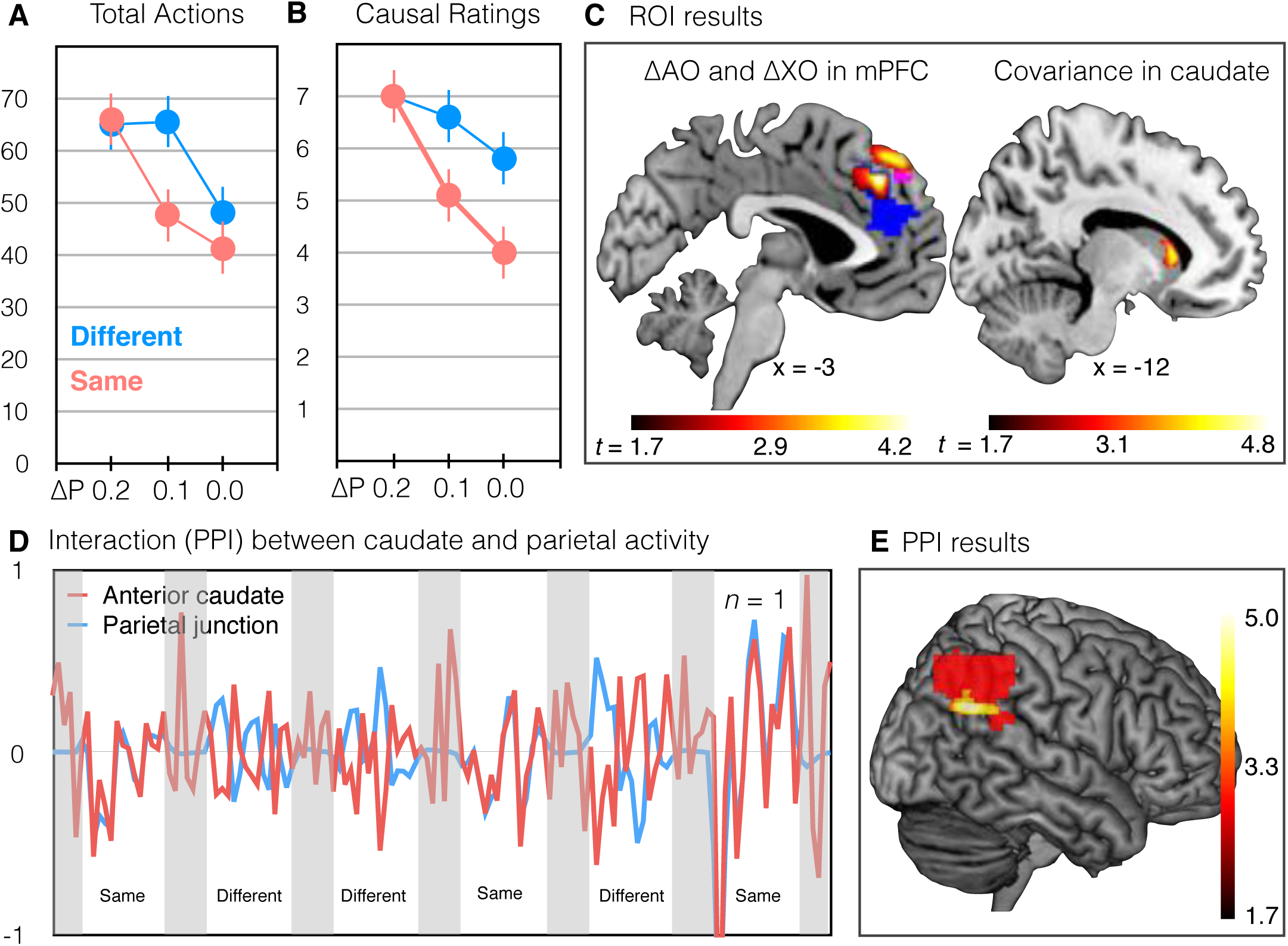
Experiment 2 results (*N* = 20) **A)** Mean (errorbar SEM) total responses were signficantly higher when the free outcomes were different from earned outcomes, outcome main effect *F*_1,19_ = 12.57, *p* = .02. This difference decreased as ΔP increased, outcome by contingency interaction *F*_1.5,38_ = 7.24, *p* = .005. **B)** Mean (errorbar SEM) causal judgments were higher when free outcomes were different from earned outcomes (*F*_1,19_ = 33.82, *p* = .6E-4) and this difference decreased as ΔP increased, interaction *F*_2,38_ = 10.07, *p* = .002. **C)** Updates to the action ΔAO occurred in the BA9 fROI (violet, image threshold *p* < .001 svc) while updates to the background ΔXO occurred in the dorsal ACC fROI (blue, image threshold *p* < .001 svc) and the covariance was tracked in the caudate fROI (green, image threshold p < .001 svc). **D)** Illustrative results from a single subject showing the caudate and posterior parietal cortex interacted with the causal condition **E)** Right parietal junction activity interacted with caudate activity when noncontingent outcomes were indistinguishable from contingent outcomes (covariance from Experiment 1 shown in red for comparison), image thresold *p* < .001 unc.

## Discussion

We sought to establish the learning rules that govern AO learning in instrumental conditioning and their neural bases. We found that the medial prefrontal cortex participates in a circuit that detects and segregates the unique causal effects of our actions from other background effects, and more importantly, that this segregation was generated by a Bayesian prediction-error, described by a Kalman filter, which uses a summed prediction-error term along with the covariance between potential causes to distinguish the unique effect of actions from background effects. Furthermore, the caudate appears to be a key point of integration of the covariance term and prediction-error; it segregates the summed prediction-error into separate update values for each causal belief. Thus, this model represents a simple, iterative Bayesian model of change, that unlike other computational-level models,^22^ provides an algorithmic account of AO learning that can be instantiated in the neural code.^23^

Many results have emphasized the critical role of the medial prefrontal cortex in AO learning, however the exact nature of this role has been unspecified. Indeed, there is a wealth of evidence that in the rat, the prelimbic region of the medial prefrontal cortex is critical for the acquisition of goal-directed actions.^2,24–29^ Computational models of medial prefrontal cortex function in humans such as the PRO model^1^ assume the *dorsal anterior cingulate* signals negative prediction-errors during AO learning, consistent with other prediction-error models.^8^ Such models only offer a partial explanation of the results we observed here: The update to the background effect (ΔXO) represents the learned probability of the noncontingent outcome that must be adjusted against the probability of the contingent outcome. This is consistent with negative prediction-errors when positive changes to the background (+ΔXO) occur in the context of a negative covariance term, and so result in negative changes to the AO contingency (–ΔAO). At other times, positive changes to the background probability (+ΔXO) will occur in line with changes to the AO contingency, consistent with positive prediction-errors. This occurs when the covariance is positive, for instance, in situations in which we cannot distinguish the unique effect of our actions from potential background causes.

Our results also distinguished a separate region of the mPFC, near the medial surface of BA9, whose activity represented updates to the unique effect of the action (ΔAO) and replicated the involvement of the mPFC in an independent sample, highlighting the reliability of the current findings (Figure 6C). We further showed using PPI (Figure 6D & E) that the caudate interacts with the parietal cortex when free outcomes are indistinguishable from the earned outcome (i.e., when there was no signal or the free outcome was the same as the earned outcome). The selective interaction between parietal cortex and caudate arises under the additional demands when there is no observable information to distinguish control. When noncontingent outcomes are indistinguishable, the covariance between our actions and the background is the only information that can be used to distinguish them. The right posterior parietal junction identified in the PPI was the same cortical region identified in Experiment 1 as tracking the covariance term (along with the caudate in the subcortex). It also overlaps with the cortical region previously implicated in learning the transition matrix during model-based reinforcement learning,^30^ suggesting, across studies and laboratories, that this region represents the covariance structure of the environment. While PPI does not indicate the direction of influence, these results are consistent with a network extending from the parietal cortex to caudate to mPFC, which tracks the covariance between actions and other events and then segregates the error-term to learn about and distinguish between the influence of different causes.

We found the competitive allocation of causal belief to actions relative to the background is a form of selective learning closely related to cue-competition models in associative learning.^8^ An important difference is the covariance matrix of the Kalman filter, in which the off-diagonal terms track the covariation between events. In our model, the covariance allowed the learner to distinguish or segregate the effects of an action from the background in the absence of that action, as shown in Figure 4. This allowed the model to reason counterfactually about what would have happened if an action had not occurred. In this manner, the covariance is analogous to heuristically motivated formalizations of within-event learning (e.g., negative alpha) which allows learning about absent events in recent versions of cue-competition models.^31,32^ Our simple Kalman filter thus combines key features of contemporary associative learning and model-based reinforcement learning.

These results also make an important contribution to the common claim that goal-directed learning is analogous to MBRL. In general, MBRL is concerned with building a model of the environment, given the state caused by each action (i.e., the covariance or transition matrix). In such models, “state prediction-errors” and the covariance matrix they update^30^ only describe the contiguity between states and, as a result, the covariance matrix cannot learn or accurately represent a causal relationship. By contrast, causal learning is not concerned with the transition probabilities between different states, but rather the trade-offs between competing contingencies to determine whether that state was cause by an action or not. This is a primary difference between causal models and MBRL. The question of which action caused which state is arguably more fundamental to goal-directed learning. For example, the prediction-errors we observed here are critically different from “state prediction-errors” because they are adjusted for the probability that another state (the background) may also cause the outcome. This adjustment leads to a representation of the unique causal strength of our actions. Hence, an important implication of this proposal is that MBRL *per se* may be sufficient for maximizing reward but it does not provide a complete account of goal-directed learning since it is unable to calculate, and so is insensitive to, the causal relationship between actions and states.

Unlike some other computational models of causal learning, the causal estimates learned by our Kalman filter converge to ΔP, a normative measure of causal strength. Other researchers have argued that ΔP does not provide the best approximation of human causal inference because changing the base-rate probability of an outcome while holding ΔP constant modulates causal judgements (e.g., “the base-rate illusion”). However the base-rate illusion is considerably weaker in free-response, instrumental learning where trials are not explicitly segmented.^33–36^ Furthermore, when learning about causal effects, active intervention is a more reliable guide to causal relations than is sheer observation, largely because actions constitute one basic way to control for possible alternative causes.^37–40^ Humans are able to reason suppositionally or counterfactually about what would be expected to happen if an intervention is made or not made, and midbrain dopamine neuron firing^41^ along with our Bayesian model reflects these counterfactual action values. For these reasons, AO learning may not suffer the same biases as other forms of causal learning that are based on passive observation.

In conclusion, learning about the causal effects of our actions, as required for goal-directed learning and as investigated here, appears to reflect features of traditional associative models such as competition for predictive value, as well as modern conventions such as environmental structure (covariance). In our hands, these features were combined in a highly simplified, iterative Kalman filter that learned a probability distribution over action-outcome contingencies to provide a novel account of AO learning. In our results there was impressive agreement across experiments and replications that distinct regions of a corticostriatal network distinguished the unique causal effect of actions from those of the background. More generally, the results revealed how our neuroanatomy performs Bayesian computations, consistent with growing evidence that the brain learns and make decisions on the basis of probability distributions.^23,42–44^

## Methods

### Participants

In Experiment 1, 31 right-handed English speaking volunteers, aged between 19 and 51 (mean age 30.5) were scanned. One participant was removed due to excessive head movement (> 2 mm), thus n = 30 (18 females). On the basis of a power analysis (Supplementary Material), scans from 23 right-handed, English speaking volunteers, aged between 17 and 32 (mean age 26) were considered for Experiment 2. Three participants were removed due to excessive head movement (> 2 mm), thus *n* = 20 (11 females). All participants were free of food allergies, neurological or psychiatric disease, and psychotropic drugs, and reported strong liking of the snack foods we provided as reward. Informed consent to participate was obtained and the study was approved by the Human Research Ethics Committee at the University of Sydney (HREC no. 12812). Participants were reimbursed $45 AUD in shopping vouchers, in addition to the snack foods they earned during the test session.

### AO contingency degradation task

In each experiment, participants were instructed not to eat three hours prior the appointment. Pre-testing involved obtaining preference ratings on a 7-point scale for each of three snacks (M&Ms, BBQ flavored crackers, chocolate cookies), from which the two most similarly preferred snacks were selected for the experiment.

Experiment 1 involved learning two AO contingencies concurrently. Participants were instructed they could liberate snack foods (BBQ flavored crackers and M&Ms) from a vending machine by tilting it to the left or right (by pressing either a left or right button), and that sometimes the vending machine would also release a snack for free. They were also instructed to find the best action for releasing snacks. Outcomes were indicated by the presentation of a visual stimulus depicting the snack for 1-s duration (a particular snack food, e.g., M&M or BBQ cracker), during which further outcomes could not be earned. The relationship between actions and outcomes were constant across blocks for each participant (e.g., left = M&M and right = BBQ crackers for all blocks). Each block lasted 120-s, and the software controlling the task PsychoPy2 v1.8, ^45,46^ divided each block into 120 one-second intervals to determine the outcome rate. Participants were unaware of the 1-s intervals, and they responded freely using the index finger on their right hand to press the left or right button on a Lumina MRI-compatible response pad (LU-400, Cedrus Corporation, CA). An action (tilt left or tilt right) earned a particular outcome with a probability *P* = 0.2 if that action had occurred in the preceding 1-s interval. If both actions occurred in the preceding interval then only the most recent action was considered for reinforcement. A free outcome was delivered with *P* = 0.2 if neither action had been made. This schedule ensured two important features: 1) that there was no serendipitous contingency between an action and a free outcome, which would result in a higher reward contingency for the contingent action^16^, and 2) the earned outcome appeared at a varying interval up to one second after a successful action, which is sufficient to introduce ambiguity into the perceived AO contingency.^47^ Participants completed six blocks; the outcome (BBQ cracker or M&M) that was subject to contingency degradation was counterbalanced across blocks (ABBAAB). At the end of each block, participants rated how causal each action was with respect to each outcome on a Likert scale from 1 (not at all) to 7 (very causal). A follow-up test was conducted after the scan. The test setting, duration and programmed AO contingencies in the follow-up test were exactly the same as in the scanner, with the addition of a 1-s yellow light cue displayed on the front of the virtual vending machine immediately prior to the delivery of each free reward. At the end of all testing, participants received all snacks that had been delivered onscreen during test.

Experiment 2 involved learning a single AO contingency. The session was arranged in 12 blocks of 60-s duration, and in each block the participant responded freely for a single snack food reward (counterbalanced between BBQ crackers and M&Ms). As before, in each block the positive contingency was *P*(O|A) = 0.2 in every second a response was made. The probability of a free outcome in every second when no response was made, i.e., *P*(O|~A), varied between 0, 0.1 and 0.2 across blocks in a counterbalanced order. Conversely, ΔP varied from 0.2, 0.1 to 0. In addition, Experiment 2 varied whether the free outcome was the same snack or a different snack as the earned outcome in each block, in an ABBA order. In half the blocks the earned and free outcomes were different which effectively signaled the free outcomes and allowed the participant to discern the causal effect of their actions. For the other half of blocks the free outcomes were the same snack food as the earned outcome, thus making it difficult to discern unique causal effects.

### Behavior data analysis

The behavioral data consisted of the rate of responding during each block and the causal ratings obtained at the end each block. Experiment 1 tested for differences between Con and Deg actions in the proportion of total responses, as well as mean causal ratings. In each case, a Shapiro-Wilk test confirmed the data did not violate the assumption of normality and differences were assessed by paired t-tests (two-tailed). Experiment 2 tested the main effect of the outcome condition (Same versus Different free outcomes) and its linear interaction with the contingency condition (ΔP = 0.2, 0.1, and 0.0), using a 2 × 3 repeated measures ANOVA (two-tailed). Mauchly’s test was used to detect violations of sphericity, in which case the Greenhouse-Geisser correction was applied.

### Model-based analysis

For each of the learning models described below, the real-time occurrence of outcomes was modelled with a logistic function f(x) = 1 / (1 + e^∞ (D-k)^) to produce a binary result (0,1) determining whether the outcome will or will not be associated with the prior action. D is the delay between the previous action and outcome, and k is the temporal threshold included as a free parameter in the model-fitting described below.

#### Prediction-error learning

The prediction-error model adopted a standard delta rule exemplified in the Rescorla-Wagner model^8^ and adapted for vector-valued predictions in modern reincarnations.^1^ This allows multiple action outcomes to be predicted simultaneously, each with its own summed error-term. In this model, the predicted outcome **Ô** is a weighted sum of actions and background cues (**Ô** = *V***a**). Updates to the weights (*V)* occur by the prediction error:

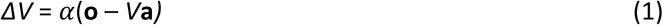

where α is a free parameter controlling the learning rate. In this way, the model replicates the prediction-error learning of the Kalman algorithm below, but without adjustment by a covariance matrix so all changes are restricted to actions on the current trial.

#### Model-Based Reinforcement Learning

The MBRL model was adapted from the FORWARD model described in Glascher et al (2010), which uses experience with state transitions to update an estimated matrix of transition probabilities. The transition matrix (*T*) held the current estimate of the probability of transitioning from action **a** (a binary vector indicating one of three possibilities: make action A1, make action A2, or Wait) to an outcome state **o** (a binary vector indicating one of three possibilities: outcome O1 delivered, outcome O2 delivered, or no outcome delivered [background state]). Wait actions occurred at the end of every second in which no other action occurred. In the *T* matrix, the different actions were represented in different columns, while the different outcomes were represented in different rows. The transitions were initialized to uniform distributions connecting each action and outcome. Upon each step, having taken action **a** and arrived in outcome state **o**, the FORWARD learner estimates the expected outcomes on the basis of the current transition matrix (**Ô** = *T***a**), and computes a state prediction-error *ΔT*:

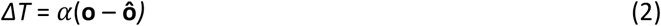

Updates to the transition matrix *T* of the observed transition occur via Δ*T*

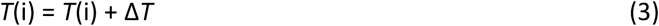

where *i* is the column corresponding to the taken action.

#### Kalman algorithm

The aim of the Kalman algorithm was to learn the unique causal weight of each action over and above the background (i.e., ΔP). The algorithm builds a probabilistic representation of the causal weights (**w**) of each input (actions and background cues) predicting each outcome (**o**), representing causal beliefs. The causal beliefs are represented by a multivariate normal density *N*(**w**|**µ***,C*) with a prior Gaussian distribution, and after observing each outcome the causal weights are updated by changes in the mean and variance of each distribution (see below). The mean **µ** represents the belief in the unique causal strength while the variance *C* captures the uncertainty around that belief. When the variance is large, there is large uncertainty regarding the true causal strength. The updating equations for the mean and variance of the causal weight have the following form:

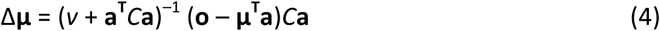

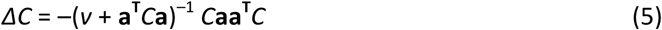

where *v* is a free parameter capturing outcome variance. For each second, the execution of an action and the constant background context are represented as a binary input vector (**a**) and an outcome vector (**o**) represents the delivery of the outcomes (O1 and O2). The first term (*v* + **a^T^***C***a**)^−1^ represents the total certainty (inverse sum of outcome uncertainty and belief uncertainty) and it governs the learning rate. **µ^T^a** is a vector representing the learned causal weights on the basis of the current inputs (**a**). The difference between the observed outcome and learned causal weights (**o** – **µ^T^a**) is a vector of outcome-specific prediction errors, each of which represents a summed error-term for O1 and O2. The rightmost term *C***a** is the product of the covariance matrix and input vector, and it allows for changes to the mean belief about actions otherwise correlated with the background context but absent on the present trial. In this manner, the Kalman filter is able to distinguish the unique influence of actions from the context during noncontingent outcomes. Importantly, the covariation between each action and the background are tracked in the off-diagonal elements of *C*, which allowed us to test a unique prediction of this model. Thus changes to the mean beliefs (Δ**µ**) depend on the prediction-error as well as the covariance matrix *C*.

#### Null model

For comparison, we also described a null model without any temporal dynamics but ideal asymptote performance. The null model assumed that the probability of taking each action was proportional to the final *ΔP* obtained in each block, so *Q*_right_ = *ΔP*_right_ and *Q*_left_ = *ΔP*_left_.

#### Policy

The policy of each model was the same. In each learning model, each action had a unique causal relationship with two outcomes representing the belief regarding that particular AO contingency (e.g., **µ**_*a,o*_). So for each action, we selected the highest causal belief associated with that action *Q*_a_ = arg max_a_ **µ**_*a,o*_ and then used it to probabilistically explain the action choices of each participant using the softmax rule:

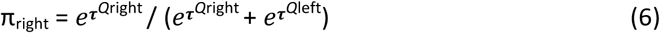

#### Bayesian model comparison

We generated observation models based on the three learning models described above as well as the null model, and fit them to each subject’s behaviour separately using maximum-likelihood estimation. A non-linear optimization was achieved using the fmincon function in MATLAB R2014B (The Mathworks Inc., MA, USA) over 100 random starting values for each subject. We measured the overall goodness of fit using the average likelihood per trial of the best fit model for each subject. The average likelihood per trial was calculated as the exponent of the sum of log likelihoods divided by the number of trials (responses) for each subject. We compared models by aggregating the probability of the data given the model over each subject’s fit (single-subject Bayesian Information Criterion [BIC] score) to estimate the model evidence for the full dataset. The aggregate for each model was then compared to compute a Group Bayes Factor (GBF). We also report the number of subjects for whom individual model comparison gave the same answer as the GBF and the positive evidence ratio (PER), where “positive” evidence for one model versus another exists if the log Bayes factor is larger than three (Kass & Raftery, 1995).

### Image data analysis

MRI data were acquired on a 3-Tesla GE Discovery using a 32-channel head coil. A T1-weighted high-resolution structural scan was acquired for each subject for screening and registration with a 1-mm^3^ voxel resolution (TR: 7200 ms, TE: 2.7 ms, 176 sagittal slices, 1-mm thick, no gap, 256 × 256 × 256 matrix). For BOLD acquisition, we acquired echo planar image (EPI) volumes comprising 52 axial slices in an ascending interleaved acquisition order (TR: 2910 ms, TE: 20 ms, FA: 90 degrees, FOV: 240 mm, matrix: 128 × 128, acceleration factor: 2, slice gap: 0.2 mm) with a voxel resolution of 1.88 × 1.88 × 2.0 mm. Slices were angled 15 degrees from AC-PC to reduce signal loss in the OFC. In Experiment 1, 343 EPIs were acquired while in Experiment 2, 260 EPIs were acquired.

Data were analysed using SPM8 (www.fil.ion.ucl.ac.uk/spm). Preprocessing and statistical analysis were conducted separately for each experiment. The first four images were automatically discarded to allow for T1 equilibrium effects, then images were slice time corrected to the middle slice and realigned with the first volume. The structural image was coregistered to the mean functional image, segmented and warped to MNI space. The warp parameters were then used to normalise the resampled functional images (2 mm^3^). Images were then smoothed with a Gaussian kernel of 8-mm full-width half maximum to improve sensitivity for group analysis.

### Model-based fMRI analysis

For each first-level GLM analyses, we constructed a vector of delta values for action causal beliefs (ΔAO) and background causal beliefs (ΔXO), generated with the parameters provided by the group maximum likelihood estimation (MLE).^48^ For ΔAO, the delta values were taken for the current action contingency (i.e., A1-O1 or A2-O2), while for ΔXO the delta values were taken for the current outcome (B-O1 or B-O2). To test for brain activity tracking the unique changes in each vector, we entered ΔAO and ΔXO as parametric modulators of a stick function that included both response and reward times in an event-related design. While these update signals will fluctuate independently, there will be some collinearity when the covariance is zero. Collinearity is a problem when trying to determine unique effects associated with each regressor. However, the variance inflation factor can be used to indicate if a collinearity problem is present. The variance inflation factor was 1.23, which is within the bounds of a conservative threshold < 5.^49^ Nevertheless, to remove any residual collinearity between these regressors, each regressor was entered as the second modulator to ensure it was adjusted for the prior regressor using the default orthogonalize routine in SPM.^49^ Each GLM also contained rating periods and six movement parameters. Betas were estimated with a 128-s high-pass filter and AR1 correction for auto-correlation. The resulting beta images were included in a group-level random effects analysis in SPM one-sample t-tests. SPM F-contrasts (two-tailed) were used to create whole-brain statistical parametric maps, corrected for multiple comparisons using a voxel-level FWE-p < .05. SPM t-contrasts (one-tailed) were used in each ROI analysis, corrected for multiple comparisons using FWE (svc) in the case of anatomical ROIs (Experiment 1) and uncorrected at *p* < .001 (svc) in the case of independent functional ROIs (Experiment 2).

#### DCM analysis

Each of the volumes-of-interest (VOI) for the DCM analysis was spatially defined according to the group results of the relevant GLM analysis. The BA9 VOI was defined by the significant cluster from the analysis of ΔAO in the group results. This significant group cluster was used to construct a binary mask and this mask was then used to define the VOI and extract the first eigenvector for all individuals, adjusted for the ΔAO and ΔXO regressors. All subthreshold voxels within the mask were included which were *p* < .5 (uncorrected), which roughly corresponds to all voxel activity positively related to ΔAO. The mPFC VOI was extracted in the same manner but using the significant cluster from the analysis of ΔXO in the group analysis. The caudate VOI was defined by a group ROI analysis of press rate restricted to the striatum (*p* < .05, small volume corrected), with a single cluster of 48 voxels in the right caudate (peak MNI: +15 +10 +6), and otherwise extracted in the same manner as other VOIs.

#### PPI analysis

The psychological term was the block condition, whether or not the free rewards were distinguishable within that block. The physiological term was the timeseries from the anterior caudate in each participant (n = 20) using a group fROI mask from the GLM analysis of the covariance. We constructed the interaction term in SPM8 (per defaults) and included all three terms in the first-level GLM. Finally we tested for regions of interaction in the whole-brain, corrected for multiple comparisons FDR -*q* < .05.

### Data availability

The analyses in this report were conducted by RWM (unblinded). Data is available upon request. Unthresholded statistical maps are available for viewing and download at http://neurovault.org/collections/VXWZKTWE/. Experimental programs and Matlab code to generate simulations can be downloaded from http://balleinelab.com

## Acknowledgments

This study was supported by a Laureate Fellowship from the Australian Research Council (ARC; #FL0992409) awarded to BWB. RWM was supported by National Health and Medical Research Council (NHMRC) Project Grant #1069487, and the ARC Centre of Excellence in Cognition and its Disorders (Macquarie University). MJG was supported by the NHMRC R.D. Wright Biomedical Career Development Fellowship (APP1061875). MLP was supported by an ARC Future Fellowship (FT100100260) and BWB by a Senior Principal Research Fellowship from the NHMRC. The programs used in the behavioral tasks are available for download at http://balleinelab.com.

The authors declare no biomedical financial interests or potential conflicts of interest.

## Supplementary Figure

**Figure S1.**
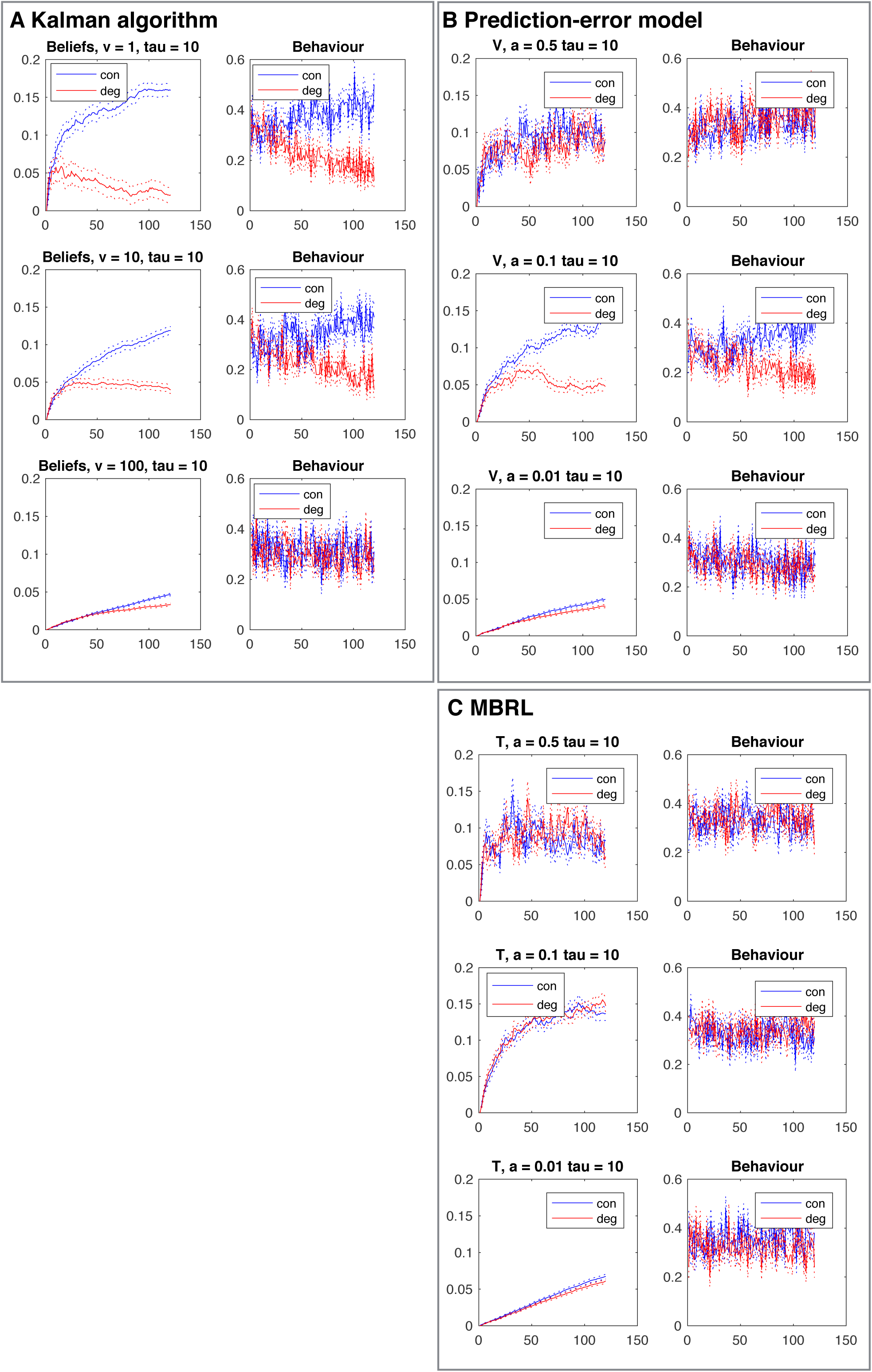
Simulation results (mean and SEM of 100 iterations) showing the temporal dynamics of learning and behaviour for each model at representative parameter values. **A)** Beliefs (**u**) and behaviour in the Kalman algorithm distinguish causal actions over time throughout parameter space **B)** *V*-values and behaviour of the PE model distinguish causal actions over time in a narrower range of paramter values **C)** State-action transitions (T) and behaviour in MBRL fails to distinguish causal actions

**Figure S2.**
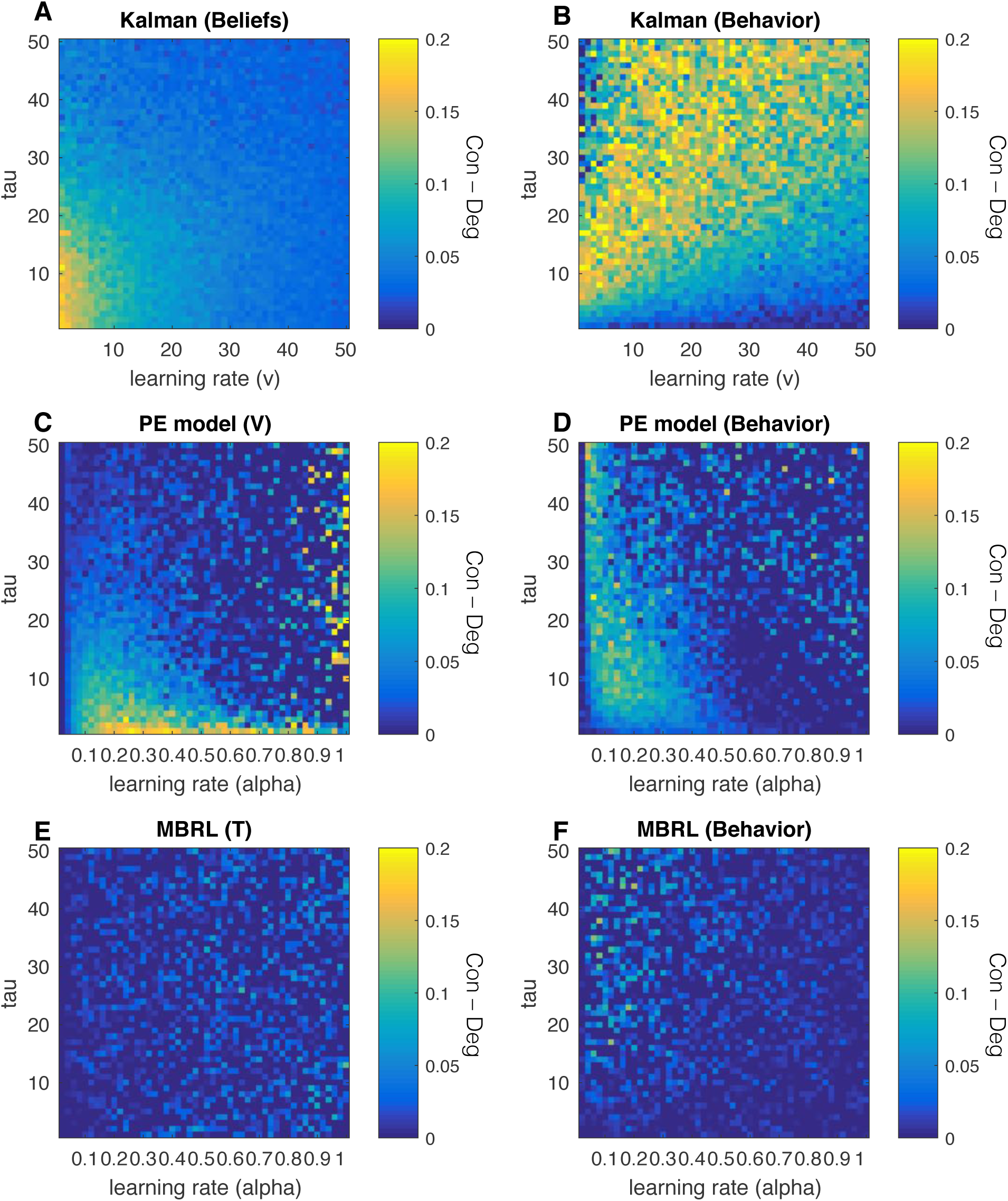
Simulation results showing the parameter space (of learning rate & tau) over which each model distinguishes the causal action during degradation. **A)** Beliefs (**u**) in the Kalman algorithm distinguish causal actions throughout parameter space, with the best distinction at low parameter values **B)** Behavior of the Kalman algorithm distinguishes causal actions throughout parameter space **C)** *V*-values in the PE model distinguish causal actions at low values of tau **D)** Behavior of the PE model partially distinguishes causal actions at low learning rates **E)** State-action transitions (T) in MBRL fail to distinguish causal actions **F)** Behavior of the MBRL fails to distinguish causal actions

**Figure S3.**
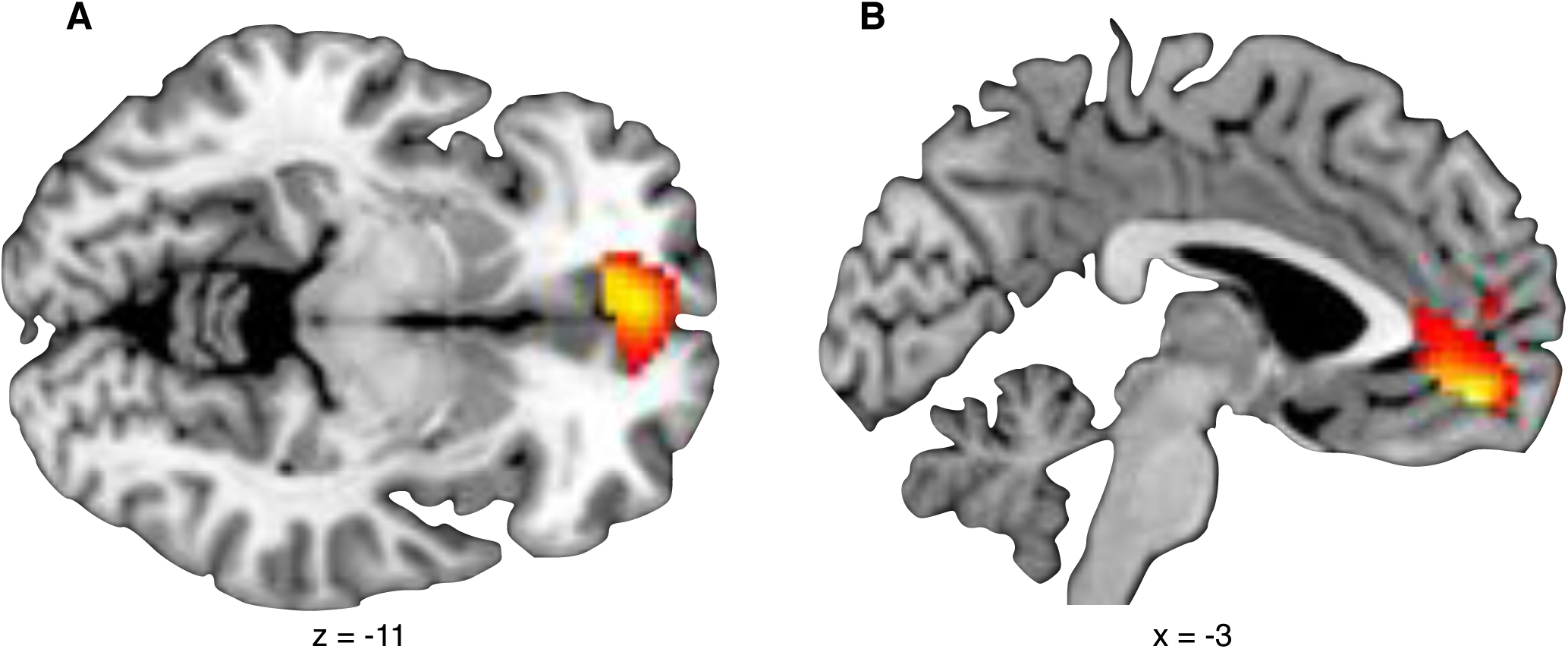
Responses to the alternate AO contingency in the ventromedial prefrontal cortex (N = 30), *F*_1,29_ = 18.08, FWE = .011, including **A)** the medial orbitofrontal cortex and **B)** the anterior cingulate. Image thresholded at p < .001, uncorrected.

**Figure S4.**
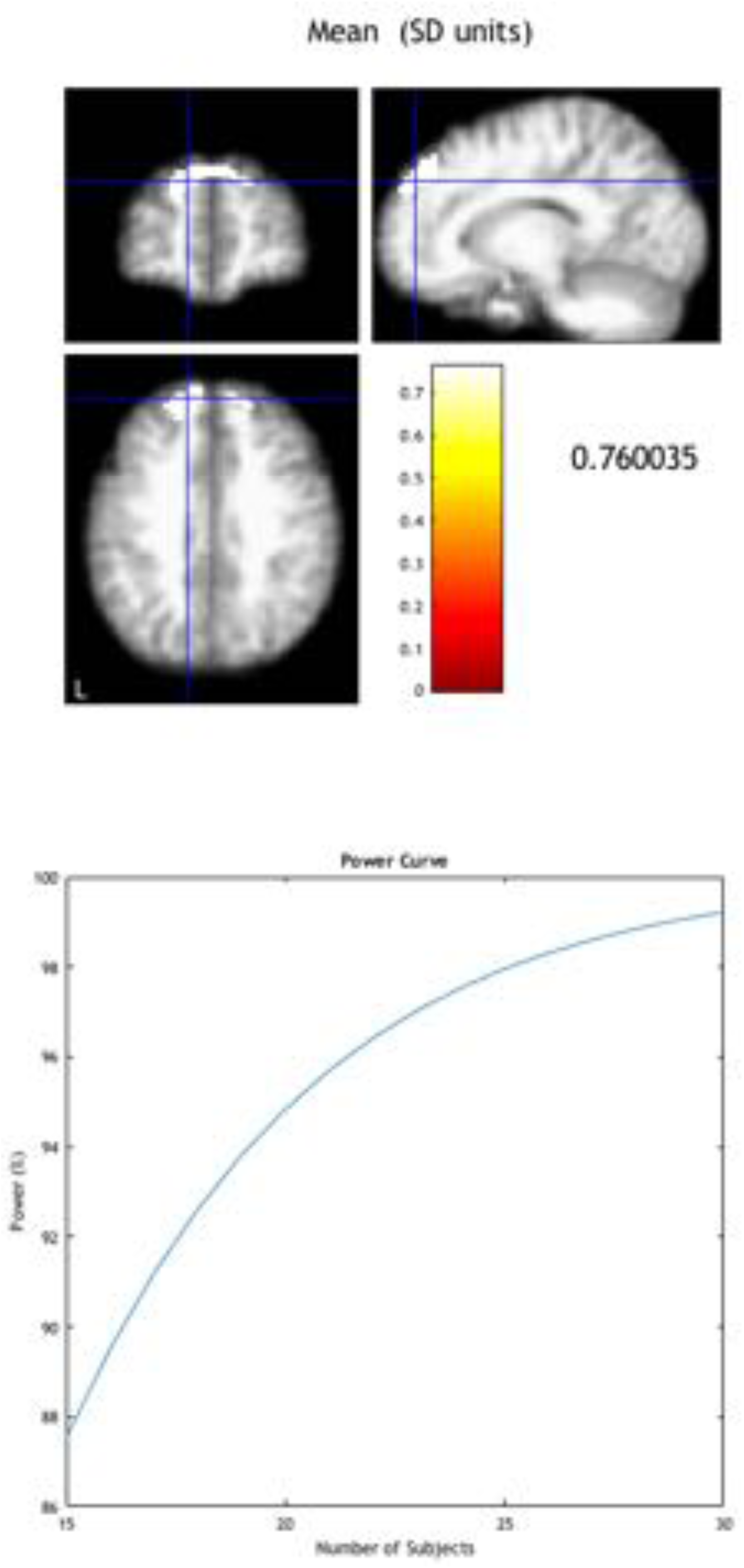
Power analysis of Experiment 1, using the mPFC response during ΔAO, indicated N > 20 would achieve 95 percent power

